# Ectomycorrhizal fungi associated with mature white spruce (*Picea glauca*) across three Alaskan treelines

**DOI:** 10.64898/2026.08.20.745949

**Authors:** Kristina Kuprina, Saroj Basnet, Manuela Bog, Martin Schnittler

## Abstract

Root-associated fungal (RAF) communities influence tree nutrition, yet their drivers and link to tree growth remain unclear, particularly near treelines. We characterized RAF communities, with a focus on ectomycorrhizal (ECM) fungi, associated with fine roots of 111 mature white spruce (*Picea glauca* [Moench] Voss) trees at three Alaskan sites (Alaska Range, Brooks Range, and Interior Alaska) across paired forest and treeline plots representing three treeline ecotones, two elevational and one drought-limited. Using ITS2 metabarcoding, we related composition, alpha diversity and guild structure to individual tree growth (raw, detrended and regionally standardized basal area increment; n = 106) averaged over 5–30 years. Our results showed that site was the strongest predictor of composition (19.3% of explained variation), followed by soil pH (11.5%), tree age (2.4%) and habitat type (2.1%). Treeline effects were small and context-dependent: 13, 17 and 2 taxa differed between habitat types in the Alaska Range, Brooks Range and Interior Alaska, respectively, independent of tree age; alpha diversity was lower at the treeline only at the drought-limited site; and ECM relative abundance was higher at all three treelines (significantly so in the Brooks Range), while plant saprotrophs declined at all three. Six of seven responding ECM taxa were enriched at the treeline. RAF composition and diversity metrics were not associated with growth over any period. Thus, RAF communities of mature *P. glauca* were structured mainly by site and soil chemistry, with the most consistent treeline signature being a shift towards ECM fungi. ECM richness alone did not predict tree growth.

## 1. Introduction

Boreal forests are shaped by strong interactions between trees and their belowground biotic environment. Among them, associations with ectomycorrhizal (ECM) fungi are particularly common, with fungal partners facilitating plant nutrient acquisition in exchange for carbon supplied by their hosts (Read and Perez-Moreno 2003; van der Heijden *et al*. 2008; Smith and Read 2010). At the same time, saprotrophic fungi play a central role in decomposing plant polymers, such as cellulose and lignin, thereby contributing substantially to carbon and nutrient cycling in forest ecosystems (Lindahl *et al*. 2007; Smith and Read 2010; Clemmensen *et al*. 2013; Ransedokken *et al*. 2024). Together, these and other fungal groups form diverse root-associated fungal (RAF) communities, whose composition can be influenced by multiple factors, including habitat type, soil properties, elevation, and even host age (Sterkenburg *et al*. 2015; Luo *et al*. 2023; Hofmann *et al*. 2023; Tedersoo *et al*. 2024; Bai *et al*. 2024; Agan *et al*. 2026). Determining the relative importance of these factors is particularly relevant in environmentally stressful habitats, where limiting climatic conditions and resource availability may influence interactions between trees and their fungal partners.

Environmental filtering is expected to be especially pronounced at treelines, where conditions become increasingly limiting for tree establishment and growth. Compared with closed-canopy forests, treelines are exposed to harsher environmental conditions, including lower temperatures, greater wind and drought stress, while soils are often shallower and less nutrient-rich, all of which can constrain tree growth (Sullivan *et al*. 2015; Müller *et al*. 2016). Thus, the treeline provides a natural setting in which to investigate whether these limitations are linked with the composition of RAF communities.

Despite numerous greenhouse experiments demonstrating that various ECM fungal species can stimulate growth of seedlings (e.g., Dixon *et al*. 1984; Hoeksema *et al*. 2010; Kipfer *et al*. 2012; Smith *et al*. 2015; Zimmermann *et al*. 2025), there is no substantial evidence whether they play a role in modulating tree growth at older life stages in the natural forest (Anthony 2025). Field-based, observational studies complement the experiments and help to understand how variation in RAF communities relates to forest functioning.

White spruce (*Picea glauca* [Moench] Voss) provides a promising system for examining these relationships. This tree species is widely distributed across boreal and subarctic forests of North America (Nienstaedt 1990). In Alaska, it is a major species at both northern and elevational treelines and occurs near the dry limit of conifer growth in Interior Alaska (Lloyd *et al*. 2013). *Picea glauca* has been the focus of numerous studies investigating tree responses and adaptation to environmental and climate variability (McGuire *et al*. 2010; Lloyd *et al*. 2013; Nicklen *et al*. 2016; Hyne*s* and Haman 2020; Zacharias *et al*. 2021, 2022). Its growth has been shown to exhibit substantial plasticity, with local microenvironmental conditions explaining more variation in growth responses than genetic similarity among trees (Zacharias *et al*. 2021). Furthermore, Zacharias *et al*. (2022) found that *P. glauca* trees growing at the treeline exhibited greater climate sensitivity in radial growth than trees in paired forest plots, whereas genetic variation explained little of the observed variation in tree growth. Together, these findings highlight the importance of local environmental conditions in shaping *P. glauca* growth and suggest that interactions with belowground biota may contribute to tree performance and responses to environmental stress.

Despite the importance of fungal associations for tree nutrition and ecosystem processes, the relationship between RAF communities and growth of *P. glauca* remains poorly understood. To our knowledge, no previous study has examined the RAF community of mature *P. glauca* trees. Here, we investigated RAF communities associated with *P. glauca* fine roots across natural ecotones in Alaska, including one drought-limited (Interior Alaska) and two cold-limited (Brooks Range and Alaska Range) treelines. We used individual metabarcoding of fine roots together with measurements of recent tree growth to examine whether RAF communities vary between paired forest and treeline plots and whether variation in fungal communities is associated with tree growth.

Specifically, we address three research questions:

i. Which factors are the strongest predictors of RAF community composition of *Picea glauca* fine roots?
ii. Do the limiting environmental conditions at the treeline affect RAF community composition?
iii. Is variation in RAF community composition and diversity associated with variation in *P. glauca* growth?

## 2. Material and Methods

### 2.1. Data collection

We studied *Picea glauca* trees at three sites in Alaska. At each site, we sampled two plots (habitat types): a nearly monodominant white spruce forest and an adjacent treeline. The Brooks Range site comprised a treeline (BT) influenced by both latitudinal and elevational gradients, situated at the northern edge of the distribution of *P. glauca* on a steep south-facing slope, with the corresponding forest plot (BF) located ca. 22 m apart and 50 m lower in elevation. The Alaska Range site represented an elevational treeline (AT) on a south-facing slope ca. 1,000 m a.s.l., with the paired forest plot (AF) situated approximately 1.3 km away and 200 m lower in elevation. The Interior Alaska site represented a drought-limited treeline (IT) located on a steep (12–34°) south-facing bluff, with the adjacent forest plot (IF) situated on a shallow slope ca. 150 m a.s.l. In total, six plots were sampled: BF and BT, IF and IT, and AF and AT (Fig. 1). More characteristics of the study plots are provided in Wilmking *et al*. (2017), Trouillier *et al*. (2018) and Zacharias *et al*. (2022).

**Figure 1.**
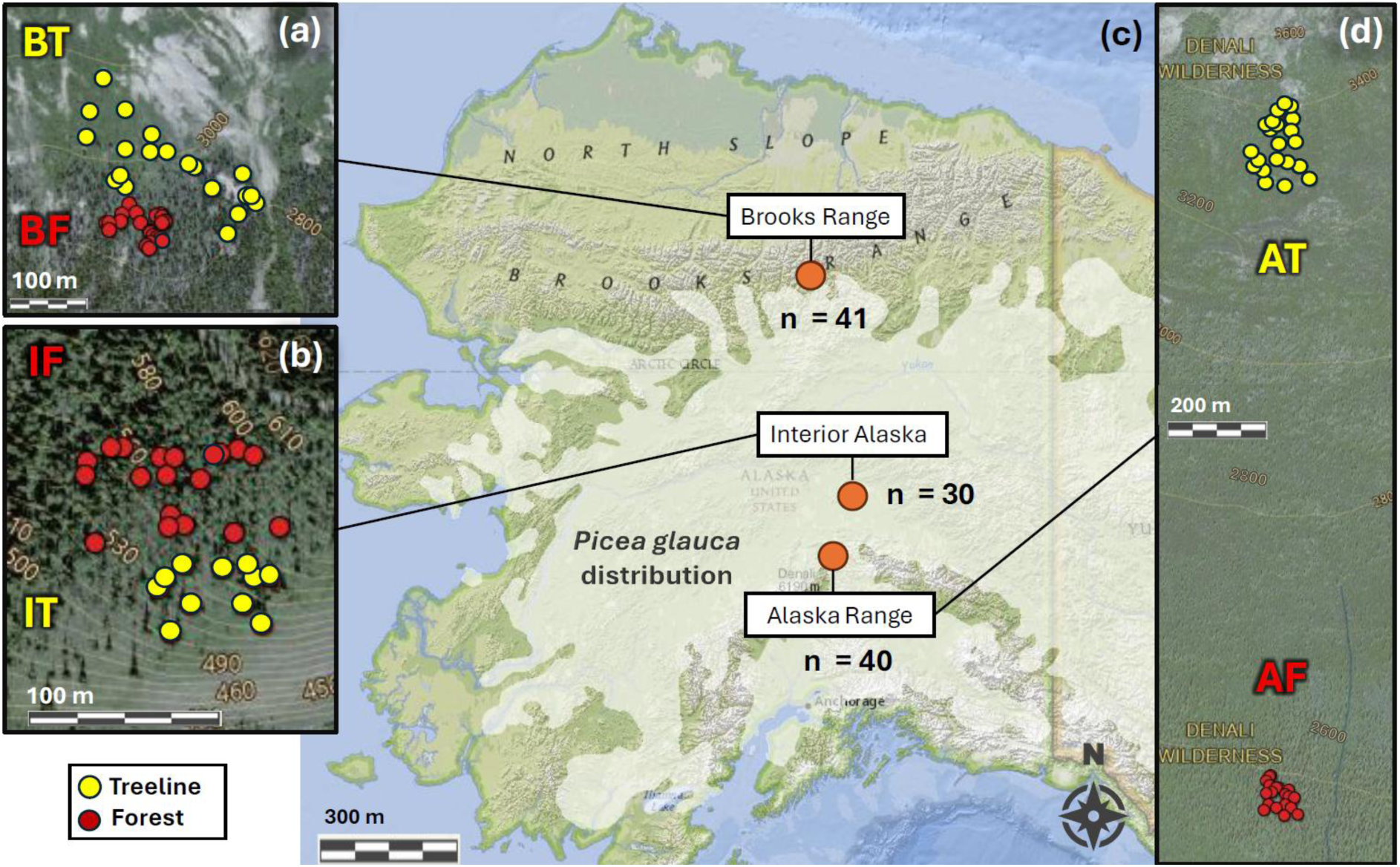
Geographic distribution of sampled *Picea glauca* trees across six plots in (a) the Brooks Range, (b) Interior Alaska and (d) the Alaska Range sites. Panel (c) displays the natural distribution of *Picea glauca* in Alaska (shaded area; modified after Little 1971). The elevation labels are in feet.

Sample collection was conducted between 20 August and 3 September 2022. We selected adult trees that appeared visually healthy and recorded position (GPS coordinates; Suppl. Tab. ST1). For each studied tree, we measured height, diameter at breast height (DBH), and extracted wood cores to estimate age and growth parameters. From each sampled tree, two increment cores were taken at breast height perpendicular to each other to reduce a possible bias due to irregular/eccentric growth. All the data were obtained for 106 trees, in total.

To assess root-associated fungi (RAF), six fine-root fragments (each at least 15 cm long) were collected from each tree, with samples taken as far apart around the trunk as possible. To ensure that each root sample belonged to the correct tree, we traced the root directly from the trunk. Each root sample was cleaned of large substrate fragments through gentle shaking and then stored in an individual plastic zip-lock bag. The root samples were air-dried at room temperature for at least two weeks.

One teaspoon of soil was collected from three spots adjacent to fine roots around the tree. The three soil samples were pooled into a single plastic zip-lock bag and then dried at room temperature for one month. The pH of soil samples was then measured with pH/mV pocket meter WTW pH330i with electrode WTW SenTix41 (both Xylem Analytics, Weilheim i. OB, Germany) according to DIN ISO 10390, after incubating 5 g of soil in 25 mL of 0.01 M CaCl_2_ for one day. In total, root and soil samples were collected from 111 trees including 52 trees from treeline plots and 59 trees from forest plots (Fig. 1).

### 2.2. Tree growth characterisation

The increment core samples were glued to wooden sample holders and were sanded and polished using a sanding machine with different grades of sandpaper (grit size of 120–600) in order to make annual rings visible. The samples were then scanned using a conventional scanner (Epson Expression 12000XL). Annual tree-ring widths (TRW) were measured on the scanned image with 0.01 mm precision using the software CooRecorder v9.3 (Cybis Elektronik and Data AB, Sweden) and all radii were cross-dated visually and using CDendro v9.3.1. For a more detailed description of all study sites, tree core sample collection and core processing see Eusemann *et al*. (2016), Wilmking *et al*. (2017) and Trouillier *et al*. (2018).

We used three measures of tree growth: basal area increment (BAI), detrended BAI (detBAI) and BAI indices from regional curve standardization (rcsBAI).

Ring-width series of the two cores per tree were averaged, and TRW were converted into BAI using the *bai.out* function of the R package *dplR* (Bunn *et al*. 2020). Starting from the stem radius derived from DBH measured at sampling, the function subtracts ring widths successively from bark to pith to reconstruct the radius at the end of each year, and calculates BAI as the difference in basal area between consecutive years. DBH was measured over bark and no bark correction was applied. One core (tree 45201) that could not be reliably cross-dated in its outer part was excluded.

DetBAI was calculated to remove long-term trends related to tree age and size. BAI series were log-transformed and detrended with a 30-year cubic smoothing spline with a 50% frequency cut-off (Cook and Peters 1981) using the *detrend* function of *dplR*. The residuals were back-transformed, so that each index expresses annual BAI relative to the tree’s own long-term trend (1 = growth on the trend). The log transformation was used because splines fitted to untransformed BAI can become negative near the pith, where BAI is close to zero (Cook and Peters 1997).

Individual detrending removes differences in growth level between trees. We therefore also calculated rcsBAI indices (Briffa and Melvin 2011), which express the growth of each tree relative to the average growth of trees of the same cambial age in the same plot. Regional curves were built from all available *P. glauca* trees of the three sites: the comprehensive treeline growth dataset from earlier sampling campaigns (2012 and 2015) plus the trees sampled in 2022, 1,279 trees in total. Separate curves were built for the forest and treeline plots of the Alaska Range (339 and 206 trees) and the Brooks Range (232 and 146 trees). For Interior Alaska, forest and treeline trees were combined in one curve (356 trees), because of the absence of a clear transition zone between treeline and forest. RcsBAI indices were calculated as the ratio of annual BAI to the regional curve value at the tree’s cambial age, so that values > 1 indicate growth faster than the reference population at the same cambial age.

As a measure of recent tree growth, we calculated mean BAI, detBAI and rcsBAI over the last 5 (2018–2022), 10 (2013–2022), 15 (2008–2022) and 30 (1993–2022) years for each tree.

We calculated differences in soil pH, tree age, height and DBH between plots. Normality and homogeneity of variances were assessed using the Shapiro–Wilk and Bartlett’s tests, respectively. As the data did not meet these assumptions, the Kruskal–Wallis test followed by Dunn’s post hoc test were conducted using the R package *rstatix* v0.7.3 (Kassambara 2023). *P*-values were adjusted for multiple comparisons using Bonferroni correction.

Data on growth parameters, soil pH, tree age, DBH and height across the study plots can be found in Suppl. Tabs. ST1-ST4. Pairwise relationships among parameters for tree growth over ten years, tree size attributes (age, DBH, height) and soil pH were assessed using Pearson correlation analysis. Analysis was conducted using the R package *Hmisc* v5.2-6 (Harrell 2026). Relationships were visualized using a scatterplot matrix generated with the extension *GGally* v2.4.0 (Schloerke *et al*. 2025) to R package *ggplot2* v4.0.2 (Wickham 2016). Linear relationships were displayed in the lower panels using ordinary least squares regression, while Pearson correlation coefficients were shown in the upper panels. Kernel density distributions of each variable were plotted along the diagonal.

### 2.3. Metabarcoding

For DNA extraction from root material, 40 fragments of visible ectomycorrhizal (ECM) root tips were picked from each root sample and pooled, resulting in 240 fragments of ECM from each studied tree.

To estimate technical error, four samples were processed in three technical replicates starting from DNA extraction. To evaluate within-tree variation in RAF community composition, we included root replicates by additionally analysing five fine-root fragments from each of three trees (one each from IF, BT and AF). Each root fragment was processed and analysed individually.

DNA extraction, amplification of ITS2 region as a barcode, DNA library preparation and Illumina MiSeq sequencing were conducted according to Kuprina *et al*. (2025). The ITS2 region was amplified using primers gITS7F (Ihrmark *et al*. 2012) and ITS4ngsR (Tedersoo *et al*. 2014). Throughout the library preparation, the samples were quasi-randomly distributed across one and a half plates. To monitor potential contamination, two PCR no-template controls (NTCs), four DNA extraction NTCs, and two Illumina sequencing NTCs were included.

In total, 142 DNA libraries were sequenced: 111 individual samples (six pooled roots each), eight technical replicates, 15 root replicates and eight NTCs.

Further data filtering and analysis was performed on the “High-Performance Computing” (HPC) cluster of the University of Greifswald, which is managed by a SLURM system and has conda v25.3.1 and python v3.7.12 installed.

Nextera adapters, as well as N and low-quality leading and trailing bases (<3, Phred-33), were trimmed using Trimmomatic v0.39 (Bolger *et al*. 2014). Quality of the raw and resulting sequences was checked by running FastQC v0.11.5 and MultiQC v1.14 tools (Andrews 2010; Ewels *et al*. 2016). Raw reads are available on NCBI (PRJNA1335163).

Obtained sequences were processed using the VSEARCH v2.30.0 tool (Rognes *et al*. 2016). The following steps were undertaken: merging of forward and reverse reads (*fastq_minovlen* = 150, *fastq_maxdiffs* = 15), quality filtering of reads (*fastq_maxee* = 0.5, *fastq_minlen* = 250, *fastq_maxns* = 0), dereplication of reads across the samples and removal of sequences with fewer than 10 copies (*minuniquesize* = 10), pooling of the samples. The VSEARCH package was also utilized for the identification and removal of chimera sequences using the UCHIME 4.2.40 (release date is 2024-07-16), applying both *de novo* and reference-based detections (Edgar *et al*. 2011). All resulting sequences with a similarity lower than 97% were clustered to the different Operational Taxonomic Units (OTUs) using “greedy” clustering in the function *cluster-size*. Each OTU was taxonomically assigned by comparing it to the UNITE v10.0 (release date 2025-02-19) with a 96% identity threshold (Abarenkov *et al*. 2025). As a result, multiple OTUs could be assigned to the same taxon. OTUs with fewer than 30 reads were excluded.

### 2.4. RAF community composition

Further statistical evaluation and visualisation of results were conducted in R 4.5.0 (R Core Team 2025) using RStudio 2023.12.1.

Prior to analysis, OTUs potentially associated with contamination were filtered out. OTUs were excluded if their total read counts in NTCs exceeded one tenth of the normalized total sample counts, calculated as total sample reads divided by the sample-to-NTC ratio (135/8). NTC samples were subsequently removed from the dataset.

The relative taxonomic composition of RAF communities among plots was visualized using bar plots generated with the R packages *microeco* v2.0.0 (Liu *et al*. 2021) and *ggplot2* v4.0.2 (Wickham 2016) based on raw read counts. In addition, the relative abundance of taxa at each site was visualized using heat trees generated with the R package *metacoder* v0.3.8 (Foster *et al*. 2017), for which raw read counts were used. Variations in taxonomic composition among different plots were visualized by conducting a Principal Coordinate Analysis (PCoA) with Euclidean distances (with and without technical replicates) for all OTUs and for ECM OTUs only utilizing the R package *phyloseq* v1.52.0 (McMurdie and Holmes 2013). Before that, the raw read counts underwent normalization for sampling depth through a variance stabilizing (VST) transformation implemented in the R package *DESeq2* v1.46 (Love *et al*. 2014).

To test whether RAF communities were spatially structured within plots, Mantel tests were performed separately for each of the six plots using the R package *vegan* v2.7-2 (Oksanen *et al*. 2025). Community distances were Euclidean distances between VST-transformed read counts (all OTUs or ECM OTUs only), as used for the PCoA, and geographic distances between trees were derived from their GPS coordinates. Three tests were run per plot: community distance against geographic distance; a partial Mantel test of the same relationship conditioned on the difference in soil pH between trees; and difference in soil pH against geographic distance. Significance was assessed with 9,999 permutations and *p*-values were adjusted across the six plots using the Benjamini–Hochberg correction.

The redundancy analyses (RDA) and variation partitioning were conducted to identify predictors of RAF community composition and assess their relative explanatory power. The *varpart* function from the R package *vegan* v2.7-2 was used. Three models were fitted: (a) “*RAF ∼ Site*”, (b) “*RAF ∼ Soil pH + Habitat Type + Condition (Site)*”, (c) “*RAF ∼ Tree Age + Condition (Site + Habitat Type + Soil pH)”*. RAF was represented by a matrix of VST-transformed read counts for each OTU per sample. Tests were performed twice: for all detected OTUs and only for ECM OTUs. Statistical significance of the models was evaluated using permutation tests (9,999 permutations, n = 106) implemented in the *anova.cca* function from the R package *vegan*.

To compare tree growth with RAF community composition, we extracted first five principal components of the Principal Component Analysis (PCA) using the *prcomp* function in R from all detected OTUs and from ECM OTUs separately. VST-transformed read counts were used. Then, we performed RDA in R package *vegan* for each time frame using the model: “*log(rcsBAI) ∼ PC1 + PC2 + PC3 + PC4 + PC5 + Condition* (*Site + Habitat Type + Soil pH*)“. In total, eight RDAs with permutation tests (9,999 permutations) were conducted. *P*-values were adjusted across the eight models with the Benjamini–Hochberg correction.

Differences in the abundance and occurrence of fungal taxa between treeline and forest trees were tested separately for each site, at genus and at species level, after summing the read counts of OTUs per taxon. Abundance was tested for taxa present in at least 30% of the trees of a site using negative binomial generalized linear models implemented in *DESeq2*, with size factors estimated by the “poscounts” method and a design including tree age (standardized) and habitat type. The habitat effect was assessed with a likelihood-ratio test against a reduced model containing tree age only, and the effect of tree age with a likelihood-ratio test against a reduced model containing habitat type only. Occurrence was tested for all taxa present in at least three trees of a site using Fisher’s exact test on presence/absence. *P*-values were adjusted with the Benjamini–Hochberg correction separately for each test, taxonomic rank and site. A taxon was considered to differ between habitats if either test was significant (adjusted *p* < 0.05); taxa are reported at genus level, or at species level where the genus itself did not differ.

### 2.5. RAF alpha diversity

To assess the relationships between tree growth and RAF diversity, we used three estimates of alpha diversity: OTU richness (number of OTUs), the Shannon diversity index and the inverse Simpson index. The estimates were calculated from raw read counts of (1) all OTUs and (2) ECM OTUs only, with the function *estimate_richness* from the R package *phyloseq* v1.52.0. Raw counts were used because OTU richness was not correlated with sequencing depth (Spearman ρ = 0.03, *p* = 0.74), and the Shannon and inverse Simpson indices are based on relative abundances.

Relationships between tree growth and RAF diversity were analysed with generalized linear models (GLMs) using the R package *mgcv* v1.9-4 (Wood 2011). The three growth measures were modelled over the last 5, 10, 15 and 30 years as follows: (i) “*BAI ∼ RAF diversity index + Habitat Type + Site + log(DBH)*” (DBH was included because raw BAI increases with tree size); (ii) “det*BAI ∼ RAF diversity index + Habitat Type + Site*”; (iii) “*rcsBAI ∼ RAF diversity index + Plot*”. In total, 72 GLMs were tested, for each growth measure and fungal dataset, 12 models were fitted (three diversity estimates × four time periods). All models used a Gamma error distribution with a log link and were fitted using restricted maximum likelihood (REML), because under the Gaussian models, residual variance increased with fitted values, and the Gamma models had lower AIC in all cases. Multicollinearity was assessed with generalized variance inflation factors (GVIF) using the R package *car* (Fox and Weisberg 2019). Adjusted GVIF values [GVIF^(1/(2×Df))] were below 1.1 for all predictors. The *p*-values were adjusted within each set using the Benjamini–Hochberg false discovery rate correction (Benjamini and Hochberg 1995). To test whether RAF diversity was related to tree size or age, DBH and tree age were modelled in the same way as a function of each diversity estimate, habitat type and site.

To visualize and test difference in OTU diversity for treelines and forests, sample-based rarefaction and extrapolation curves were generated for Hill numbers (OTU richness, Shannon, and Simpson diversities) on OTU presence/absence matrix using the R package *iNEXT* v.3.0.2 (Hsieh *et al*. 2025). Two values were considered to be significantly different when their 95% confidence intervals are not overlapping. Additionally, the package was employed to construct sample completeness curves to assess the adequacy of the sample size.

To test whether ECM diversity changes with tree age, the three ECM diversity estimates, and the relative abundance of ECM OTUs (raw reads) were modelled as a function of tree age, habitat and site, again using GLMs. OTU richness was log-transformed and modelled with a Gaussian distribution, the Shannon index with a Gaussian distribution on the original scale, the inverted Simpson index with a Gamma distribution and log link, and ECM relative abundance with a beta distribution and logit link.

### 2.6. RAF guild composition

FUNGuild v1.1 database was implemented to assign genera and species to guilds (Nguyen *et al*. 2016). If an OTU was assigned to multiple guilds, including “Ectomycorrhizal”, it was categorized as “Ectomycorrhizal”. Guilds were assigned to OTUs when confidence levels were “highly probable”, “probable” and “possible”.

The relative abundance of each guild was calculated per tree from raw read counts. Differences in the relative abundance of ECM and saprotrophic fungi between treeline and forest trees were tested separately for each site using two-sided Wilcoxon rank-sum tests in the R package *rstatix* v0.7.3 (Kassambara 2023). For each combination of guild, comparison and count type, *p*-values of the three site-level tests were adjusted using the Benjamini–Hochberg false discovery rate (FDR) correction (Benjamini and Hochberg 1995).

To test whether guild abundance was related to tree growth, we fitted GLMs using the *mgcv* v1.9-4 R package. Following the model selection procedure described in Section 2.5, we fitted the same 48 GLMs (two guilds, three growth measures, four time periods, two datasets), replacing the “*RAF diversity index”* with the relative abundance of ECM or saprotrophic OTUs as the predictor. To test whether the relationship between ECM relative abundance and tree growth differed among sites, we compared the model for mean rcsBAI over 10 years (“rcs*BAI ∼ ECM + Habitat type + Site + log(DBH)”*; Gamma distribution, log link) with a model that additionally included an “*ECM × Site”* interaction, using an F-test.

To test whether relative abundance of ECM OTUs (raw reads) changed with tree age, the GLM as a function of tree age, habitat and site, was fitted. ECM relative abundance was modelled with a beta distribution and logit link.

The scripts of the data analysis, raw data and MultiQC report are available on GitHub: https://github.com/kuprinak/Picea_glauca_ITS2.

## 3. Results

### 3.1. Data set

Soil pH ranged from 3.1 to 7.4 across all study plots (mean ± SD: 6.2 ± 1.1 in the Brooks Range, 5.5 ± 0.4 in Interior Alaska, and 3.6 ± 0.3 in Alaska Range). Brooks Range forest exhibited the widest variation in soil pH, ranging from 3.7 to 7.2 (Fig. 2a). Geographically, soil pH values in BF were found to be patchy rather than following a clear gradient (Suppl. Fig. SF1). Soil pH itself was not spatially structured within plots (Mantel test; *r* = −0.201 to 0.053, all adjusted *p* ≥ 0.76). A significant difference in soil pH was found between BF and BT plots (BF < BT; adjusted *p* = 0.0193).

**Figure 2.**
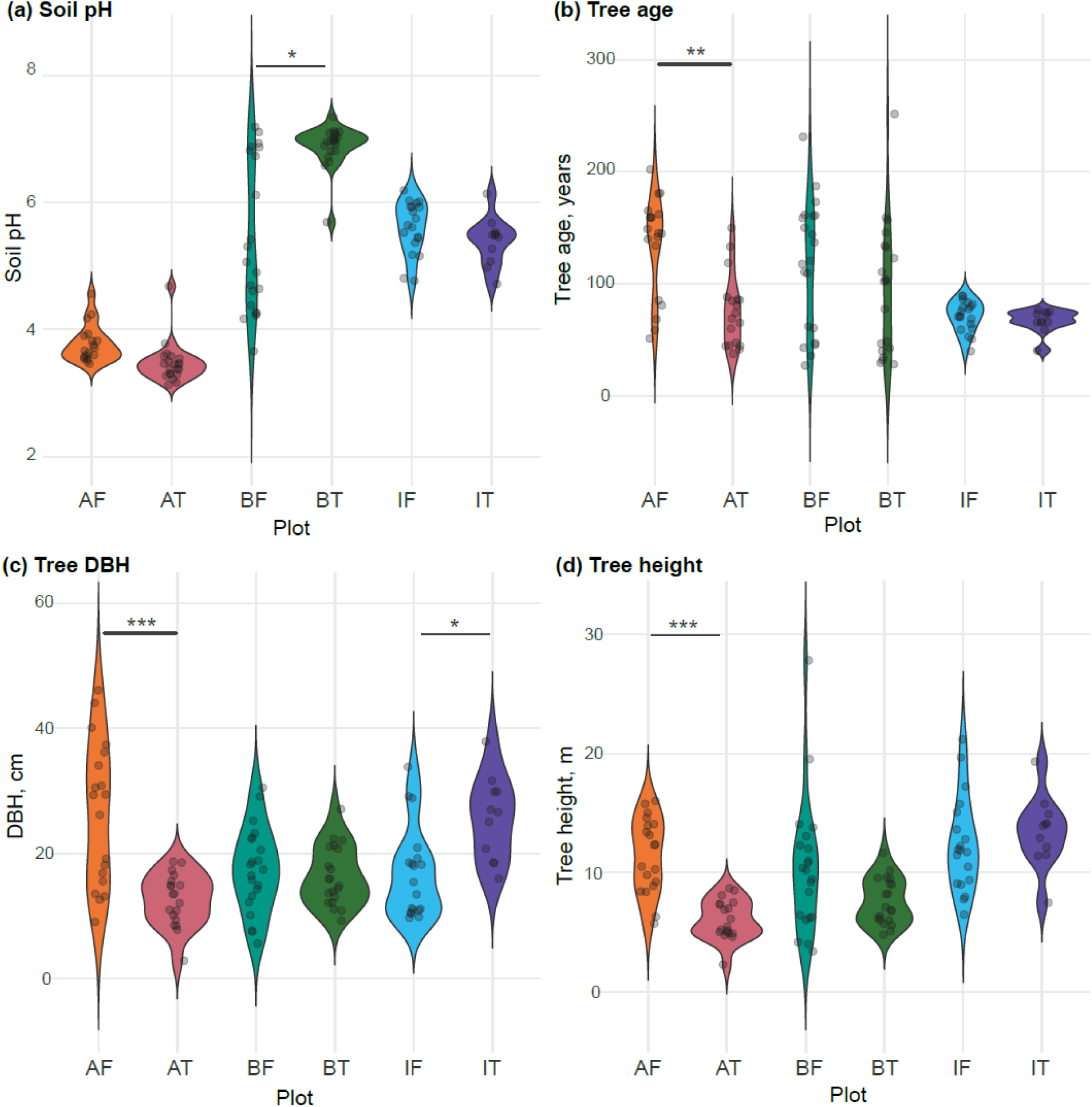
Variation in (a) soil pH, (b) tree age, (c) diameter at breast height (DBH), and (d) tree height among study plots. Significance of the difference between plots was calculated using Dunn’s test with Bonferroni *p* correction. Only significant differences between plots within a site are marked (adjusted *p*: * <0.05, ** <0.01, *** <0.001, **** <0.0001). Plots: AF – Alaska Range forest, AT – Alaska Range treeline, BF – Brooks Range forest, BT – Brooks Range treeline, IF – Interior Alaska forest, IT –Interior Alaska treeline.

Tree age ranged from 27 to 252 years. Dunn’s tests for the trees within sites showed that trees in AF were significantly older, taller and thicker than those in AT (adjusted *p* = 0.0070, < 0.0001 and 0.0002, respectively; Figs. 2b-d). Additionally, trees in IT had a larger DBH than those in IF (adjusted *p* = 0.0405; Fig. 2c).

Mean BAI over the last 5, 10, 15 and 30 years was positively correlated with DBH (Pearson *r* = 0.49–0.62, *p* < 0.001) and tree height (*r* = 0.55–0.60, *p* < 0.001) (Fig. 3). It was weakly negatively correlated with tree age for the 5- to 15-year periods (*r* = −0.21 to −0.24, *p* = 0.014–0.032), but not for the 30-year period (*r* = −0.16, *p* = 0.11). BAI was not correlated with soil pH (*r* = 0.04–0.07, *p* ≥ 0.45).

**Figure 3.**
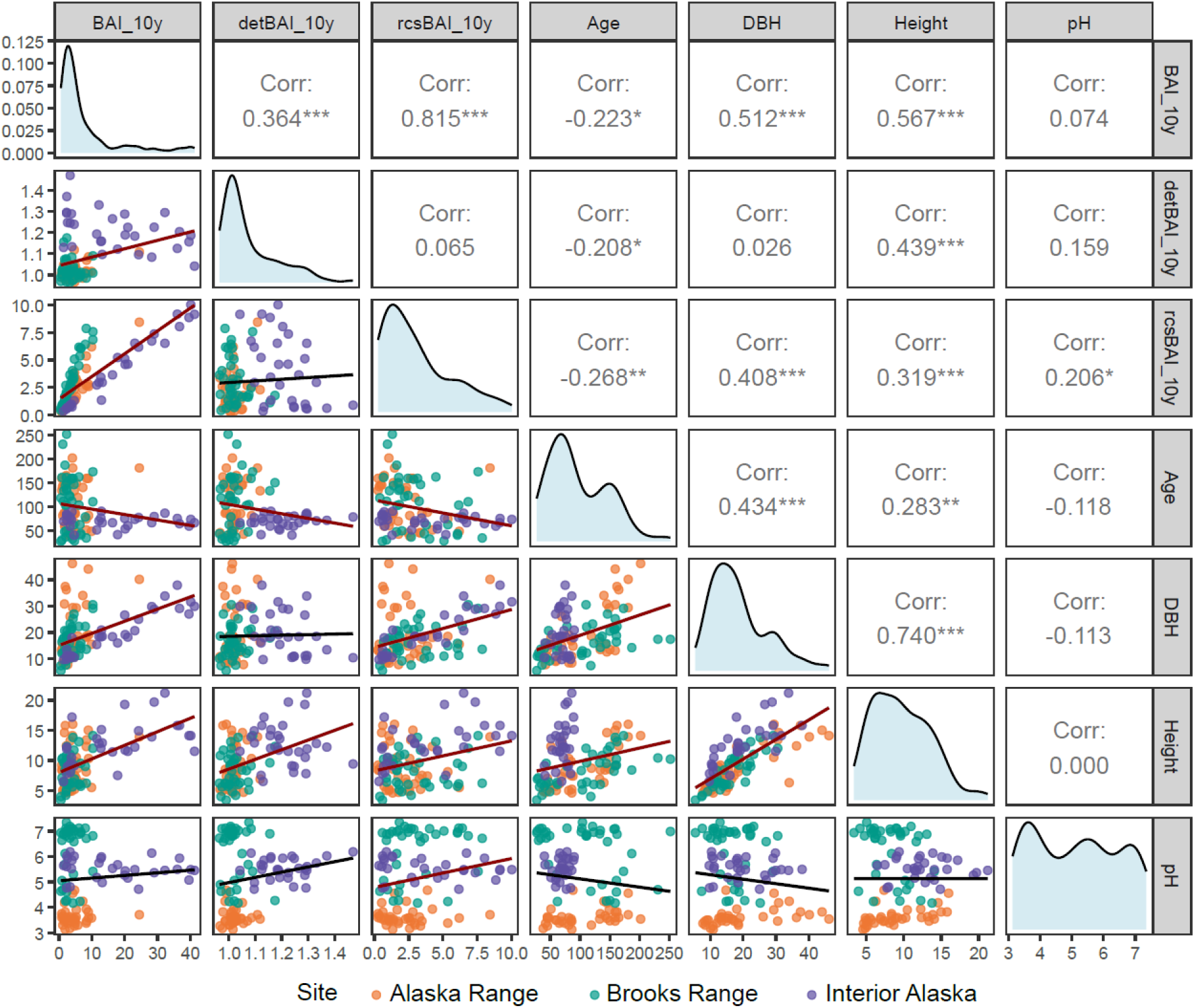
Pairwise correlations for 106 trees of *Picea glauca:* mean basal area increment over 10 years (BAI_10y), mean detrended BAI over 10 years (detBAI_10y), mean BAI after regional curve standardization over 10 years (rcsBAI_10y), age (years), dimeter at breast height (DBH, cm), height (m), and adjacent soil pH. Linear relationships are shown in the lower panels with regression fits (lines). Sample points are coloured by site. Upper panels display Pearson correlation coefficients between two variables, while Kernel density distributions of each variable are plotted along the diagonal. * *p* ≤ 0.05, ** *p* ≤ 0.01, *** *p* ≤ 0.001.

DetBAI indices showed weak and inconsistent correlations with tree age and DBH (|*r*| ≤ 0.23). Exceptions were positive correlations with tree height over 10 and 15 years (*r* = 0.44 and 0.35, *p* < 0.001) and with soil pH over 15 years (*r* = 0.40, *p* < 0.001). RcsBAI indices decreased with tree age (*r* = −0.23 to −0.28, *p* = 0.017) and increased with DBH (*r* = 0.34–0.41, *p* < 0.001).

Sample sequencing produced on average 91,124 read pairs (SD: ± 22,636 read pairs). After processing reads with Trimmomatic, 89,512 read pairs per sample remained (SD: ± 22,323 read pairs). NTCs produced on average 1,249 read pairs (SD: ± 1,273 read pairs). After processing the data with the VSEARCH tool, 692 distinct OTUs were initially identified and 467 of them were assigned to fungal taxa. Four OTUs were excluded due to potential contamination, reducing the total number to 463 OTUs.

### 3.2. RAF community composition

Taxonomic assignment successfully classified 98.7% of OTUs to the phylum, 96.4% to the class, 95.5% to the order, 92.3% to the family, 86.5% to the genus and 51.6% to the species level. In total, five phyla, 15 classes, 34 orders, 68 families, 104 genera and 212 fungal species were detected (Suppl. Figs. SF3, SF4). Basidiomycota and Ascomycota accounted for 71.9% and 22.3% of all OTUs, respectively, and for 76.4% and 21.3% of all reads in the dataset. The class Agaricomycetes was the most abundant, representing 78.3% of raw reads assigned.

Among the ECM fungi, *Cortinarius* and *Piloderma* were the most abundant genera in the Alaska Range, whereas *Tomentella*, *Inocybe*, and *Sebacina* dominated in the Brooks Range. In Interior Alaska, the ECM community was dominated by *Tomentella*, *Piloderma*, *Cortinarius*, and *Calonarius* (Suppl. Fig. SF4).

All three sites shared 76% of OTUs (351 of 463). PCoA indicated differentiation of RAF communities among sites (Fig. 4), with samples from BF being the only group overlapping with another site (Alaska Range) and showing the highest variation in both analyses, based on both the full OTU dataset and ECM-only OTUs. PCoA which included technical replicates and samples of different roots from one tree, showed higher difference between roots of one tree than between technical replicates (Suppl. Fig. SF2), supporting the use of multiple fine roots to characterize the RAF community of an individual tree.

**Figure 4.**
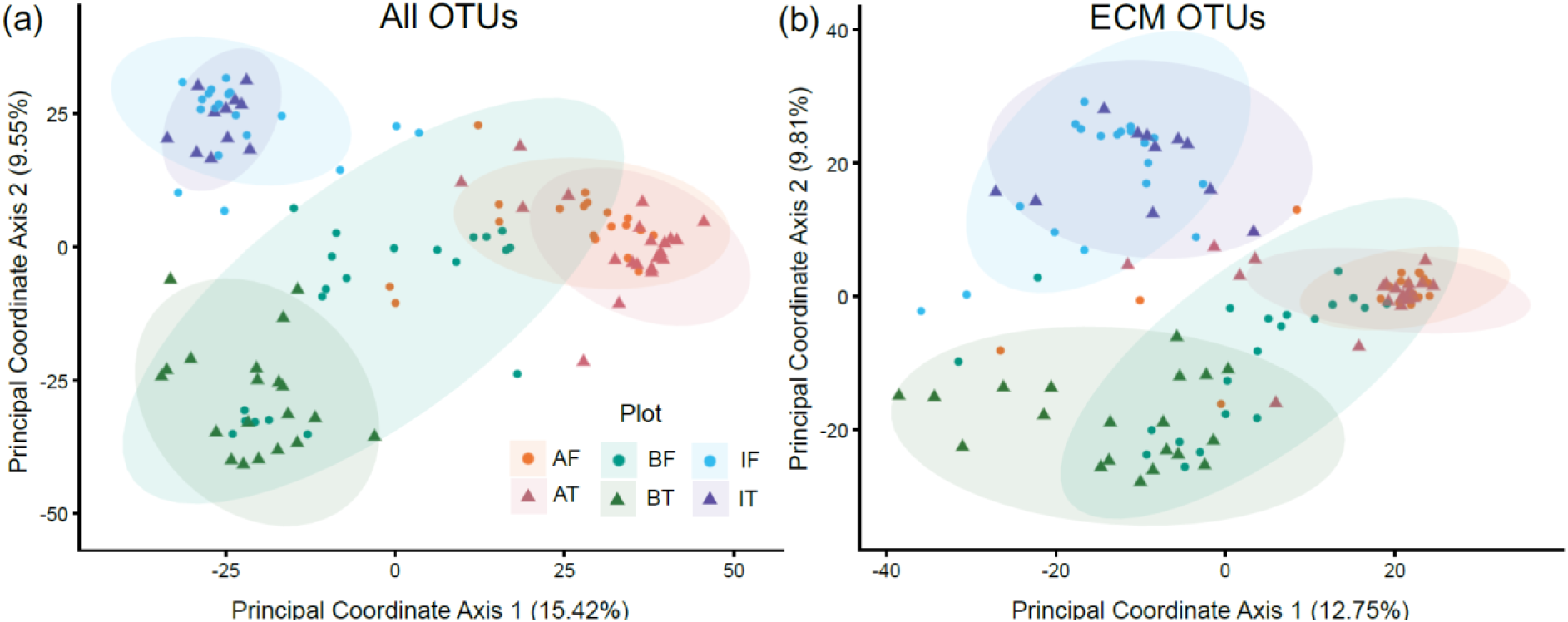
Principal Coordinate Analysis based on Euclidean distances, with 95% data ellipses illustrating within-plot clustering in ordination space, performed on six pooled root samples from 111 *Picea glauca* trees. Distances were calculated from VST-normalized counts of (a) all 463 OTUs and (b) 246 ectomycorrhizal (ECM) fungal OTUs. AF – Alaska Range forest, AT – Alaska Range treeline, BF – Brooks Range forest, BT – Brooks Range treeline, IF – Interior Alaska forest, IT – Interior Alaska treeline.

Within plots, RAF community dissimilarity was unrelated to the distance between trees in all six plots (Mantel tests: *r* = −0.127 to 0.195, all adjusted *p* ≥ 0.34), and the result was unchanged when the difference in soil pH between trees was held constant (partial Mantel tests: *r* = −0.137 to 0.199, all adjusted *p* ≥ 0.38). The same tests restricted to ECM OTUs gave equally non-significant results (*r* = −0.092 to 0.156, all adjusted *p* ≥ 0.40).

Variation partitioning analysis explained 23.2% of the variation in RAF community composition based on all OTUs (Fig. 5a) and 17.7% of the variation in ECM community composition based on ECM OTUs only (Fig. 5b). Site accounted for the largest proportion of the explained variation (19.3% and 13.9% for all OTUs and ECM OTUs, respectively), followed by soil pH (11.5% and 7.8%), tree age (2.4% and 2.0%), and habitat type, which explained the smallest proportion of variation (2.1% and 1.5%). Permutation tests for all RDA models were significant, confirming that site, soil pH, habitat type, and tree age each had a significant association with RAF community composition (all *p* < 0.05).

**Figure 5.**
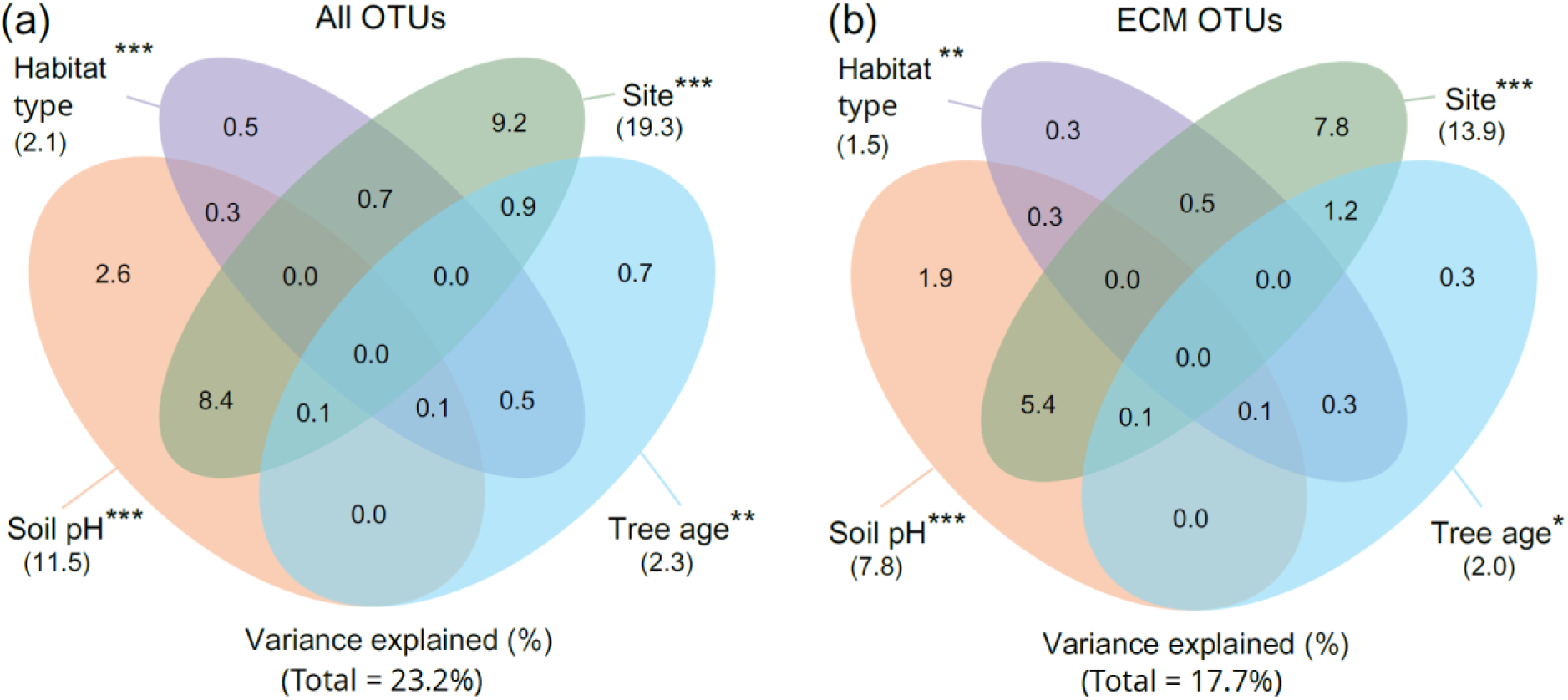
Variance in fungal community composition associated with fine roots of 106 *Picea glauca* trees explained by four predictors derived from redundancy analysis (RDA) variance partitioning. The analysed fungal community comprised either (a) all 463 detected OTUs or (b) only the 246 ECM OTUs. The statistical significance was evaluated using permutation tests for RDA models: \**p* < 0.05, \*\**p* ≤ 0.01, \*\*\**p* ≤ 0.001.

Ectomycorrhizal OTUs were the dominant trophic guild, comprising 53.1% of all OTUs and 69.0% of all reads (Fig. 6b). Saprotrophic OTUs accounted for 29.8% of OTUs and 21.1% of reads.

**Figure 6.**
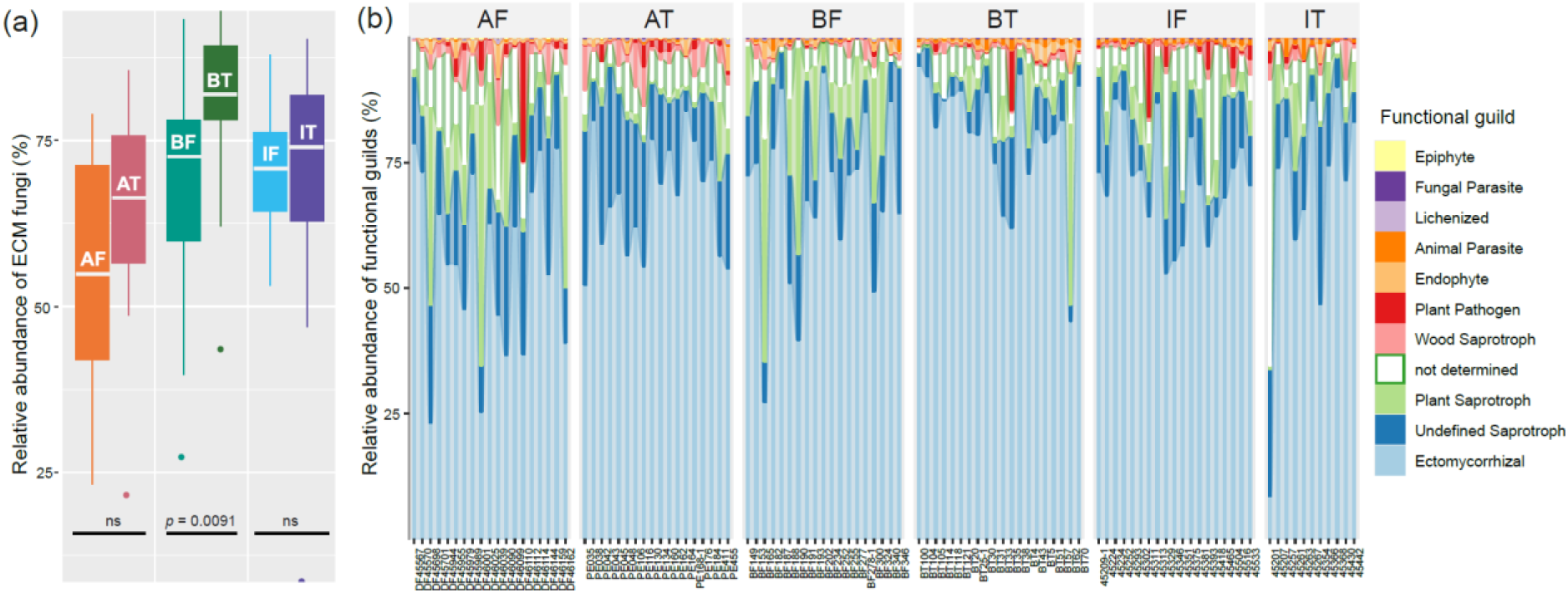
Relative abundance of (a) 246 OTUs assigned to ectomycorrhizal (ECM) functional guild and (b) all 463 OTUs assigned to different functional guilds and associated with fine roots of 111 *Picea glauca* trees across six study plots. AF – Alaska Range forest, AT – Alaska Range treeline, BF – Brooks Range forest, BT – Brooks Range treeline, IF – Interior Alaska forest, IT – Interior Alaska treeline.

### 3.3. Habitat type

PCoA did not reveal visible differentiation of treeline RAF communities from forest communities inside the sites for all detected OTUs or only ECM OTUs (Fig. 4).

Differential abundance and occurrence analyses identified 13 taxa that differed between the treeline and the forest plot in the Alaska Range, 17 in the Brooks Range and two in Interior Alaska (Table 1). In the Alaska Range, nine taxa were more abundant or more frequent at the treeline, including the ECM fungi *Polyozellus* (11.1% vs. 2.8% of reads) and *Piloderma sphaerosporum* (5.3% vs. 0.08%; detected in 14 treeline and three forest trees), together with saprotrophs and yeasts (*Penicillium zhuangii*, *P. penicillioides*, *Mortierella basiparvispora*, *Trechispora*, *Tausonia*), the entomopathogen *Beauveria*, and *Epicoccum*, which was more frequent but not more abundant at the treeline. Four taxa were more abundant in the forest: *Sistotrema* (9.8% vs. 0.07%), *Entomortierella*, *Cryptocoryneum* and *Kockovaella*.

**Table 1.** Fungal taxa whose abundance or occurrence in fine roots of *Picea glauca* differed between the treeline (T) and forest (F) plot of a site. Abundance was tested with negative binomial models including tree age as a covariate, on taxa present in at least 30% of the trees of a site. Occurrence was tested with Fisher’s exact test on presence/absence of the taxa. Relative abundance is the mean percentage of all fungal reads of a tree; N trees is the number of trees in which the taxon was detected (Alaska Range: 19 F, 21 T; Brooks Range: 21 F, 20 T; Interior Alaska: 19 F, 11 T). “–” - no assignment. ECM – ectomycorrhizal, SD – standard deviation, n.s. – not significant.

| Site | More abundant | Taxon | Guild | N trees in F/T | Relative abundance in F (% ± SD) | Relative abundance in T (% ± SD) | Adjusted *p* abund | Adjusted *p* occur |
| --- | --- | --- | --- | --- | --- | --- | --- | --- |
| Alaska Range | F | *Cryptocoryneum* | Wood Saprotroph | 17/3 | 0.051 ± 0.046 | 0.002 ± 0.005 | 0.0019 | 0.0008 |
|  |  | *Entomortierella* | - | 18/12 | 0.351 ± 0.532 | 0.021 ± 0.028 | 0.0020 | n.s. |
|  |  | *Kockovaella* | Epiphyte | 16/9 | 0.030 ± 0.051 | 0.006 ± 0.012 | 0.0089 | n.s. |
|  |  | *Sistotrema* | Plant Saprotroph | 11/7 | 9.766 ± 14.654 | 0.068 ± 0.189 | 0.0089 | n.s. |
|  | T | *Polyozellus* | ECM | 19/18 | 2.833 ± 4.013 | 11.07 ± 11.34 | 0.0002 | n.s. |
|  |  | *Penicillium zhuangii* | Undefined Saprotroph | 15/18 | 0.023 ± 0.025 | 0.153 ± 0.167 | 0.0004 | n.s. |
|  |  | *Trechispora* | Wood Saprotroph | 13/16 | 0.032 ± 0.047 | 0.218 ± 0.255 | 0.0019 | n.s. |
|  |  | *Piloderma sphaerosporum* | ECM | 3/14 | 0.076 ± 0.310 | 5.315 ± 7.783 | 0.0029 | 0.0161 |
|  |  | *Mortierella basiparvispora* | Undefined Saprotroph | 15/18 | 0.044 ± 0.068 | 0.344 ± 0.642 | 0.0111 | n.s. |
|  |  | *Penicillium penicillioides* | Undefined Saprotroph | 12/17 | 0.020 ± 0.029 | 0.062 ± 0.072 | 0.0113 | n.s. |
|  |  | *Tausonia* | Undefined Saprotroph | 12/18 | 0.015 ± 0.020 | 0.140 ± 0.244 | 0.0171 | n.s. |
|  |  | *Epicoccum\** | Plant Pathogen | 4/14 | 0.095 ± 0.396 | 0.025 ± 0.029 | n.s. | 0.0376 |
| Brooks Range | F | *Capronia* | Undefined Saprotroph | 21/18 | 1.310 ± 1.413 | 0.212 ± 0.295 | 0.0002 | n.s. |
|  |  | *Mycena metata* | Plant Saprotroph | 14/8 | 3.714 ± 8.205 | 0.016 ± 0.036 | 0.0015 | n.s. |
|  |  | *Umbelopsis ramanniana* | Undefined Saprotroph | 16/9 | 0.101 ± 0.173 | 0.009 ± 0.014 | 0.0052 | n.s. |
|  |  | *Cladophialophora* | Undefined Saprotroph | 21/20 | 6.365 ± 5.842 | 2.941 ± 2.487 | 0.0063 | n.s. |
|  |  | *Chromolaenicola* | Wood Saprotroph | 10/3 | 0.071 ± 0.253 | 0.001 ± 0.001 | 0.0135 | n.s. |
|  |  | *Cryptocoryneum* | Wood Saprotroph | 10/3 | 0.048 ± 0.140 | 0.002 ± 0.005 | 0.0151 | n.s. |
|  |  | *Hygrophorus* | ECM | 16/13 | 8.035 ± 18.247 | 1.646 ± 6.445 | 0.0156 | n.s. |
|  |  | *Niesslia* | Plant Saprotroph | 16/11 | 0.035 ± 0.041 | 0.007 ± 0.009 | 0.0156 | n.s. |
|  | T | *Ramoconidiophora\*\** | - | 7/16 | 0.007 ± 0.015 | 0.043 ± 0.049 | 0.0028 | n.s. |
|  |  | *Tomentella* | ECM | 21/20 | 6.920 ± 6.237 | 20.66 ± 17.52 | 0.0028 | n.s. |
|  |  | *Morinia* | Undefined Saprotroph | 5/14 | 0.004 ± 0.011 | 0.059 ± 0.149 | 0.0029 | n.s. |
|  |  | *Epicoccum* | Plant Pathogen | 17/18 | 0.109 ± 0.204 | 0.736 ± 2.613 | 0.0070 | n.s. |
|  |  | *Inocybe* | ECM | 17/20 | 6.430 ± 9.579 | 18.93 ± 14.59 | 0.0469 | n.s. |
|  |  | *Mycenella* | Plant Saprotroph | 6/15 | 0.069 ± 0.121 | 0.228 ± 0.437 | 0.0474 | n.s. |
|  |  | *Sebacina epigaea* | ECM | 9/10 | 0.249 ± 1.026 | 2.148 ± 6.375 | 0.0485 | n.s. |
|  |  | *Lyomyces* | Wood Saprotroph | 0/11 | 0.000 ± 0.000 | 0.241 ± 0.552 | – | 0.0046 |
| Interior Alaska | T | *Inocybe flocculosa* | ECM | 4/6 | 0.058 ± 0.176 | 0.722 ± 2.138 | 0.0119 | n.s. |
|  |  | *Penicillium* | Undefined Saprotroph | 19/11 | 2.942 ± 3.249 | 6.090 ± 5.599 | 0.0340 | n.s. |

\* Relative abundance in F is inflated by a single tree; The only taxon that showed a significant effect of tree age (adj *p* = 0.012)

In the Brooks Range, nine taxa were more abundant or more frequent at the treeline, among them the ECM genera *Tomentella* (20.7% vs. 6.9%) and *Inocybe* (18.9% vs. 6.4%) and the ECM species *Sebacina epigaea*, together with *Exophiala*, *Ramoconidiophora*, *Morinia*, *Epicoccum* and *Mycenella*; *Lyomyces* occurred in 11 treeline trees but in none of the forest trees. Eight taxa were more abundant in the forest, including the ECM genus *Hygrophorus* (8.0% vs. 1.6%), the black yeasts *Capronia* (1.3% vs. 0.2%) and *Cladophialophora* (6.4% vs. 2.9%), *Mycena metata* (3.7% vs. 0.02%), *Umbelopsis ramanniana*, *Chromolaenicola*, *Cryptocoryneum* and *Niesslia*. In Interior Alaska, the ECM species *Inocybe ffocculosa* and the genus *Penicillium* were more abundant at the treeline.

Overall, six of the seven ECM taxa that differed between habitats were more abundant at the treeline. Tree age, included as a covariate in all abundance models, was significantly associated with only one taxon (*Ramoconidiophora*, Brooks Range; adjusted *p* = 0.012) and with none in the Alaska Range (all adjusted *p* ≥ 0.52).

Rarefaction and extrapolation curves of the three Hill numbers (OTU richness, Shannon index and Simpson index) showed no significant difference in alpha diversity between forest and treeline trees in the Alaska Range or the Brooks Range, although values tended to be higher at the treeline in both sites (Fig. 7). In Interior Alaska, by contrast, forest trees harboured significantly higher diversity than treeline trees, for all three indices and for both the full and the ECM-only dataset. Among sites, both Brooks Range plots showed significantly higher diversity than the plots of the other sites. Sample coverage exceeded 95% in all groups, indicating that the number of trees sampled was sufficient to capture most of the fungal diversity present within a plot (Suppl. Fig. SF5).

**Figure 7.**
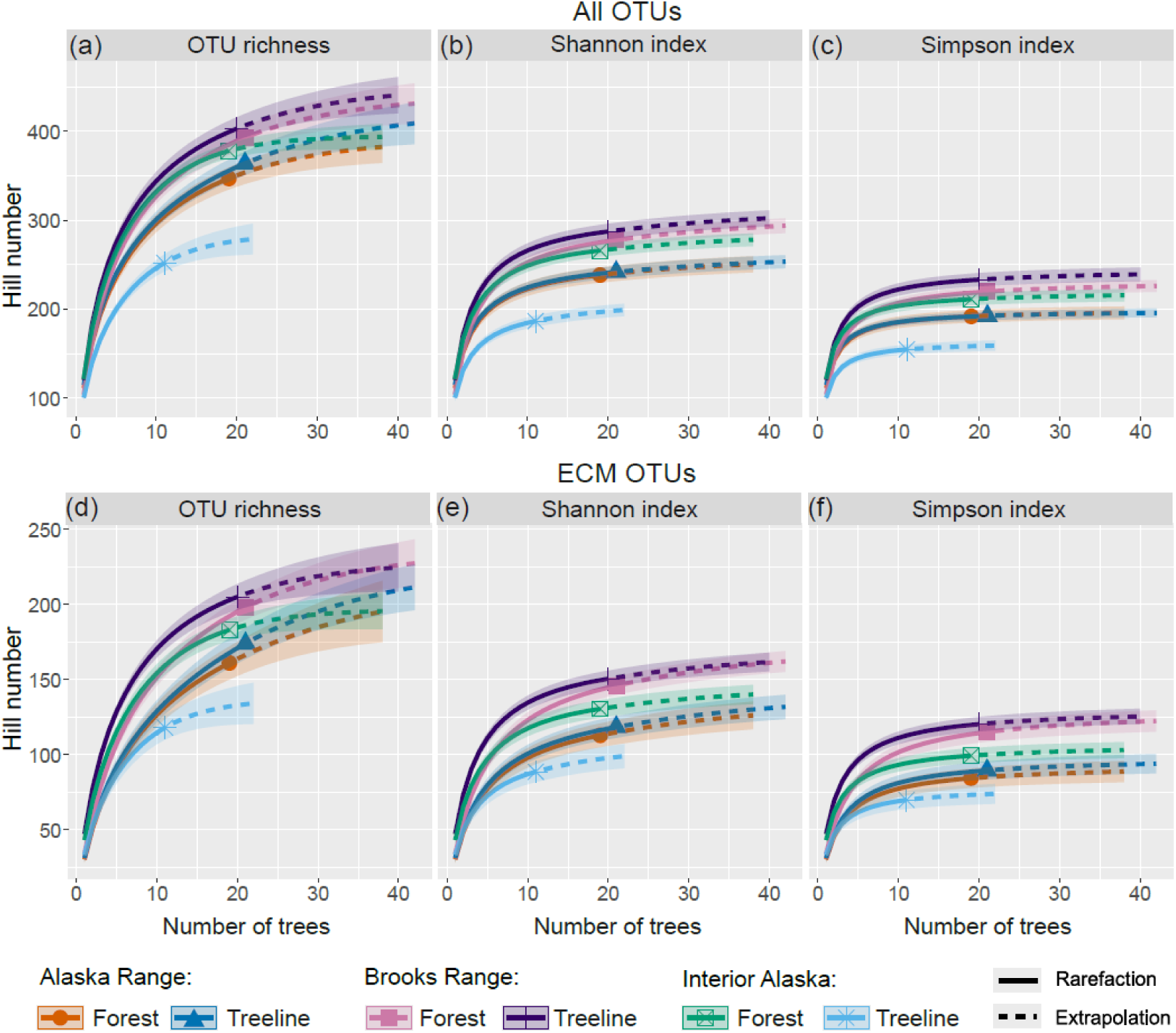
Rarefaction and extrapolation curves with 95% confidence intervals of three Hill numbers describing diversity of fungi associated with fine roots of forest and treeline trees of *Picea glauca* across three study sites: (a, d) OTU richness, (b, e) Shannon index, (c, f) Simpson index. The numbers are calculated using presence/absence data on (a, b, c) all 463 OTUs or (d, e, f) 246 ectomycorrhizal (ECM) OTUs.

The relative abundance of ECM OTUs was higher in the treelines than in the corresponding forest plots in all three sites, but significant only in the Brooks Range (median 82.0% vs. 72.6% of reads; adjusted *p* = 0.027; Fig. 6a). In contrast, the relative abundance of plant saprotrophs was lower in treelines than in forests at all three sites, significantly in the Alaska Range (1.6% vs. 4.5%; adjusted *p* = 0.032) and marginally in the Brooks Range (1.7% vs. 6.8%; adjusted *p* = 0.064). Undefined saprotrophs were less abundant at the treeline only in the Brooks Range (4.8% vs. 11.8%; adjusted *p* = 0.002), whereas wood saprotrophs did not differ between treeline and forest at any site (all adjusted *p* = 1.00).

### 3.4. RAF community and tree growth

No significant relationship was detected between RAF community composition and tree growth. All RDA models based on community ordination axes (PC1–PC5) and growth values explained no significant proportion of variation in tree growth for either the complete RAF dataset or the ECM-only dataset after controlling for site, habitat type, and soil pH (permutation tests, all *p* > 0.05).

Across all fitted GLMs, each alpha diversity index was not a significant predictor of BAI, detBAI or rcsBAI over each of four time periods (72 models; all adjusted *p* ≥ 0.416; Suppl. Tab. ST5).

None of the fitted GLMs detected a significant relationship between tree growth parameters and the relative abundance of the ectomycorrhizal or saprotrophic OTUs across sites (24 models, all *p* ≥ 0.20). The relationship between ECM relative abundance and mean rcsBAI over 10 years did not differ among sites (“*ECM × Site”* interaction: *F* = 0.52, df = 2, *p* = 0.59). None of the site-specific slopes was significant (Alaska Range: *p* = 0.53; Brooks Range: *p* = 0.52; Interior Alaska: *p* = 0.40).

### 3.5. RAF community and tree age

ECM diversity and relative abundance were not significantly related to tree age, but OTU richness and the Shannon index tended to decrease, whereas the inverse Simpson index did not change (Suppl. Tab. ST6). Neither OTU richness (estimate −0.118 ± 0.092, *p* = 0.21), the Shannon index (−0.069 ± 0.097, *p* = 0.48) nor the inverted Simpson index (0.003 ± 0.094, *p* = 0.98) changed with log-transformed tree age, and the models explained only 6–8% of the variation (adjusted R²). The relative abundance of ECM reads tended to decrease with tree age (−0.24 ± 0.14 on the logit scale, *p* = 0.10).

## 4. Discussion

### 4.1. Drivers of RAF community composition

Our results consistently identified sampling site as the strongest predictor of RAF community composition. Variance partitioning attributed 19.3% of the explained variation to site, whereas habitat type accounted for only 2.1%. The three sites differ in climate, elevation, soil chemistry and stand age, and these factors cannot be separated in our design. Interior Alaska is climatically the most distinct, with the highest mean annual temperature (−1.9 °C, compared with −8.2 °C in the Brooks Range and −3.6 °C in the Alaska Range; Zacharias *et al*. 2022) and by far the lowest elevation (ca. 150 m a.s.l. versus > 800 m at the other sites), and its trees are the youngest.

Soil pH was the second strongest predictor (11.5%), but most of this contribution was shared with site (8.9%): mean soil pH differed more among sites (3.6 in the Alaska Range, 5.5 in Interior Alaska, 6.2 in the Brooks Range) than between any pair of forest and treeline plots. The “site” effect is therefore to a large extent an edaphic effect, and our design cannot resolve how much of the residual is attributable to climate, elevation or dispersal history. Soil pH governs the solubility and plant availability of mineral nutrients and cations and thereby shapes both fungal and plant communities, directly and indirectly, and has repeatedly been identified as a dominant driver of fungal community composition in forest soils (Högberg *et al*. 2006; Glassman *et al*. 2017; Nguyen *et al*. 2020; Tedersoo *et al*. 2020; Haq *et al*. 2024; Liu *et al*. 2025; Agan *et al*. 2026). Notably, the pH gradient captured here (3.1–7.4) spans almost the full range over which boreal ECM communities turn over, and the Brooks Range forest plot alone spans 3.7–7.2, and without any accompanying difference in climate or host species. That plot also showed the greatest within-plot dispersion in the PCoA ordination, which is consistent with pH, rather than region, driving the fine-scale structure of these communities. This within-plot variation was not spatially structured (Mantel test, section 3.1), so it does not reflect the distance among neighbouring trees. Because soil pH was the only edaphic variable measured (it was chosen as a rapid, robust field proxy known to correlate with many other soil properties) the mechanisms behind this association (N and P availability, base saturation, organic matter quality) remain untested here.

Despite the strong site signal, the taxonomic composition of RAF was broadly similar across sites: 76% of OTUs were shared among all three. Spatial variation therefore reflects shifts in the relative abundance of a largely shared regional species pool rather than turnover of fungal lineages, which argues against strong dispersal limitation at this scale and points instead to local environmental sorting of a widely distributed boreal mycobiota.

Tree age also explained a small but significant fraction of compositional variation (2.4% for all OTUs, 2.0% for ECM OTUs), slightly more than habitat type. Age-related shifts in RAF communities have been reported repeatedly, with turnover of dominant taxa through stand development (Rudawska *et al*. 2018; Bai *et al*. 2024; Zhao *et al*. 2025). We detected no significant effect of tree age on alpha diversity, although OTU richness and the Shannon index both tended to decline with age (estimates −0.118 ± 0.092 and −0.069 ± 0.097 on log age), while the inverse Simpson index did not change (0.003 ± 0.094); the models explained only 6–8% of the variation. The weak negative tendency in richness contrasts with studies reporting increasing RAF diversity with stand or host age (Kyaschenko *et al*. 2017; Schön *et al*. 2023; Bai *et al*. 2024; Mandolini *et al*. 2024; Zhao *et al*. 2025), but those studies are not directly comparable: they sampled soil rather than colonized fine roots, or contrasted juveniles with mature trees. Agan *et al*. (2026) described a non-monotonic relationship between time since fire and both ECM diversity and ECM relative abundance in *Pinus sylvestris* stands, with an apparent intermediate maximum around 40–71 years since fire — although in that study neither ECM diversity nor time since fire reached statistical significance, so the pattern should be treated as a trend rather than an established relationship. Most of our trees are considerably older than that window (27–252 years, median well above 70), so a post-maximum decline remains a plausible but untested explanation for the weak negative tendency we observed.

### 4.2. The treeline signal

The forest–treeline transition left only a weak imprint on overall community structure. PCoA showed no visible separation of treeline from forest samples within any site, and variance partitioning attributed only 2.1% of the explained variation to habitat type, most of it shared with site and soil pH. Habitat type was nonetheless statistically significant, which illustrates that with 111 individual trees even a very small compositional effect is detectable.

Differential abundance and occurrence analyses were more sensitive and revealed responses at all three sites, although of very different magnitude: 13 taxa differed between forest and treeline in the Alaska Range, 17 in the Brooks Range and only two in Interior Alaska (Table 1). Most responding taxa were non-mycorrhizal. The plant saprotrophs *Mycena metata* and *Niesslia* were more abundant in the forest, consistent with the guild-level decline of plant saprotrophs at the treeline discussed below, whereas soil saprotrophs and yeasts (*Penicillium* spp., *Mortierella basiparvispora*, *Tausonia*) and the entomopathogen *Beauveria* were enriched at the treeline. Wood saprotrophs and Chaetothyriales showed opposing genus-level responses between forest and treeline, indicating no consistent guild-level response.

Among ECM fungi, by contrast, the direction of change was strikingly consistent. Six of the seven ECM taxa that differed between habitats were more abundant at the treeline: *Polyozellus* and *Piloderma sphaerosporum* in the Alaska Range, *Tomentella*, *Inocybe* and *Sebacina epigaea* in the Brooks Range, and *Inocybe flocculosa* in Interior Alaska. Several shifts were large, with *Tomentella* and *Inocybe* each roughly tripling their share of reads at the Brooks Range treeline. The only ECM taxon enriched in a forest plot was *Hygrophorus* in the Brooks Range. The treeline-enriched ECM taxa belong to four unrelated orders (Thelephorales, Atheliales, Agaricales and Sebacinales) and range from short-distance (*Inocybe*) to medium-distance fringe exploration types (*Piloderma*; Agerer 2001; Tedersoo and Smith 2013), so the enrichment does not point to a particular lineage or foraging strategy. Rather, it mirrors the guild-level result and suggests that the treeline environment favours ECM fungi broadly.

The clearest and most consistent treeline signal in our data emerged at the level of functional guilds. The relative abundance of ECM reads was higher at the treeline than in the paired forest at all three sites, reaching significance in the Brooks Range (median 82.0% vs. 72.6%), while the relative abundance of plant saprotrophs was lower at all three treelines, significantly so in the Alaska Range (1.6% vs. 4.5%) and marginally in the Brooks Range (1.7% vs. 6.8%); undefined saprotrophs were also reduced at the Brooks Range treeline (4.8% vs. 11.8%). A reciprocal ECM–saprotroph shift of this kind is expected where litter input declines and nutrient cycling become increasingly dependent on mycorrhizal mining of organic matter. Treeline stands are sparser and produce less litter than closed forest, reducing the substrate base for plant saprotrophs, while ECM mycelium and saprotrophs compete for the same organic substrates in boreal soils (Lindahl *et al*. 2007; Clemmensen *et al*. 2013; Bödeker *et al*. 2016). Barbi *et al*. (2025) similarly found saprotroph diversity to decline with elevation across 17 European elevational transects while ECM diversity peaked at mid-elevations. Sullivan *et al*. (2015) concluded that soil nutrient availability, rather than temperature directly, is the proximate constraint on treeline tree growth in northwest Alaska; an increased ECM share of the root-associated community is exactly what that nutrient-limitation hypothesis predicts. Because our root samples were obtained by picking visible ECM root tips, the absolute values of ECM relative abundance are inflated by design, but the forest–treeline *contrast* was generated with identical sampling in both habitats and is therefore interpretable.

Our result parallels Saona *et al*. (2025), who sampled the rhizosphere of *Nothofagus pumilio* at the treeline and 200 m below it at four locations spanning about 10° of latitude in southern South America across four seasons and found a marked dominance of ECM fungi at the treeline despite large variation in alpha and beta diversity among locations and seasons. That two treeline systems on different continents, with unrelated hosts and contrasting climates, converge on the same guild shift suggests it may be a general property of treeline ecotones.

Alpha diversity showed limited and site-specific responses. Significant differences in Hill numbers occurred only in Interior Alaska, where forest trees harboured higher total RAF and ECM diversity than treeline trees. Interior Alaska is our only drought-limited treeline, and moisture rather than temperature is the most likely mechanism: dry, steep, south-facing bluff soils limit hyphal exploration, spore germination and mycelial persistence, and water limitation reduces the photosynthate supply available for belowground allocation, so fewer fungal partners can be maintained per tree. This interpretation is consistent with the fact that growth of *P. glauca* in Interior Alaska is itself moisture-limited (Lloyd *et al*. 2013; Hynes and Hamann 2020; Zacharias *et al*. 2022).

Finally, that the Brooks Range was both the only site with a significant forest–treeline difference in soil pH (BF < BT) and the only site with a significant ECM abundance shift is suggestive, but with three sites the association cannot be tested and should be presented as a hypothesis.

Taken together, treeline effects in this system are real but modest and expressed through different axes at different sites: the abundance of individual taxa (strongly in the Alaska and Brooks Ranges, weakly in Interior Alaska, where the treeline plot comprised only 11 trees), alpha diversity (Interior Alaska only), and (most consistently) the balance between ECM fungi and saprotrophs, visible both in guild-level relative abundance at all three sites and in the direction of nearly all ECM taxa that responded.

### 4.3. Tree growth and RAF community

We found no evidence that RAF communities track individual tree growth. RDAs relating the first five community axes to rcsBAI over 5, 10, 15 and 30 years were non-significant for both the full and the ECM-only datasets after conditioning on site, habitat type and soil pH; none of the 72 diversity models and none of the 24 guild-abundance models reached significance; and the ECM × Site interaction was far from significant.

The effect sizes make these results an informative null rather than an absence of evidence. Across all 72 models, a one-standard-deviation increase in fungal diversity was associated with changes in growth ranging from −8.1% to +8.7%, with 95% confidence intervals spanning no wider than −21.9% to +26.4% (Suppl. Tab. ST5). For detrended BAI, which removes the age-related trend in each tree, the intervals were substantially narrower, with all effects bounded between −1.0% and +3.5%.

Our finding is consistent with the existing evidence on ECM richness. Anthony (2025) reviewed 30 studies and found that ECM richness stimulated tree growth in only 11 of them (37%), with no link detected in the majority, and concluded that differences in ECM community composition and functional identity are more informative predictors of tree growth than richness. Consistent with that, Anthony *et al*. (2022) linked ECM composition and functional genes (not richness) to a threefold difference in tree growth rate across 15 European countries, and Ibáñez *et al*. (2025) found that associations between mycorrhizal taxon abundances and the growth of mature trees were non-linear and contingent on soil nitrogen availability. Context dependence may itself explain the negative result. Jonsson *et al*. (2001) found that ECM richness increased *Betula pendula* productivity on low-fertility substrate but reduced *Pinus sylvestris* productivity on high-fertility substrate, with no effect in the reciprocal combinations.

Several non-exclusive explanations for the absence of a relationship deserve consideration.

Functional redundancy. Boreal ECM communities are species-rich but may be functionally redundant for the traits that matter to the host, so that taxonomic richness is largely decoupled from the rate of nutrient return per unit carbon. Under this view, what matters is which partners dominate and how much they deliver, not how many are present.

Growth may not be the trait that responds. At cold-limited treelines, radial growth is widely considered sink-limited (constrained by the temperature dependence of cell division and wall thickening) rather than limited by the supply of carbon or nutrients (Körner 2012). Improved nutrient acquisition would then be expressed as needle nitrogen content, root or mycelial biomass, reserve accumulation or reproductive output rather than as BAI.

A dominance-based mechanism has been proposed for seedlings: Livne-Luzon *et al*. (2017) showed that homogenizing soil inoculum led to dominance by a single, highly beneficial ECM taxon, lower overall ECM diversity and improved *Pinus halepensis* performance, whereas point inoculation allowed several taxa to proliferate and produced higher diversity but poorer performance. Hamberg *et al*. (2024) likewise found that Norway spruce saplings propagated from fast-growing parent trees hosted fewer resource-demanding ECM fungi and shorter roots, while saplings from slow-growing parents carried more Ascomycete taxa and free-living saprotrophs. Our data do not support an equivalent mechanism in mature *P. glauca*: the inverse Simpson index, which is sensitive precisely to dominance, was unrelated to growth in every model, and the relative abundance of the ECM guild as a whole was likewise unrelated. If a dominance mechanism operates here, it must act through the identity and physiological activity of the dominant partner rather than through the evenness of the community, and testing it requires quantification of colonization, nutrient transfer and taxon-specific function rather than sequence-based community descriptors.

### 4.4. Limitations and outlook

Our study has several limitations that should be considered. First, we characterized the tree–fungus relationship through community composition and diversity, and did not quantify ECM colonization intensity, extraradical mycelial biomass, root biomass or root morphology. Root function and nutrient-return rates, which are more plausibly linked to host growth than taxonomic composition (Agerer 2001), therefore remain unmeasured.

Second, our fungal data represent a single sampling occasion in late summer, whereas growth is integrated over decades; seasonal and interannual turnover in ECM communities cannot be assessed from this design.

Third, the observational design cannot establish the direction of causality. Experimental manipulation of fungal communities, combined with measurements of ECM colonization, root function, carbon allocation, nutrient availability and growth, and ideally repeated sampling across seasons and years, would be required to determine whether fungal community properties influence tree performance or reflect it.

Within a nine-year project RESPONSE, we studied growth of *P. glauca* with the goal to identify environmental drivers to explain growth patterns. Age of trees and performance, measured as DBH or BAI, was much more loosely correlated than expected, and the performance of trees within a plot differs tremendously due to shifting climatic regimes (Ohse *et al*. 2012). Microsatellite markers (Eusemann *et al*. 2014) and dendrochronological methods (Buras *et al*. 2016, Trouillier *et al*. 2019) were established in parallel, but we did not find a significant effect of certain genotypes (Zacharias *et al*. 2021); neither we could find a convincing association with single genes (Zacharias *et al*. 2022). Growth variation within clones was comparable to that within genetically different cohorts (Wilmking *et al*. 2017), and the extent of clonality in this species seems to be driven by environment (Würth *et al*. 2018). Differences in communities of endophytic fungi, reflecting potential parasite load, were as well mainly caused by environment and phenology (Eusemann *et al*. 2016, Würth *et al*. 2019). This study added ECM fungi as a potential factor but found again no single drivers of performance. As a conclusion, we deal with an extremely complex system exhibiting high phenotypic plasticity (Pampuch *et al*. 2021, Zacharias et al. 2025) in response to fluctuating climate (Lange et al. 2020), where a plethora of different factors may cause minute annual differences in growth which accumulate (likely reinforced by intraspecific competition and drought stress, Zacharias *et al*. 2022) over the years to tremendous differences in individual performance of trees.

## 5. Conclusions

Our results show that RAF communities associated with fine roots of mature *Picea glauca* in Alaska are structured predominantly by site and soil chemistry. Sampling site and soil pH together accounted for most of the explained variation in community composition, and their contributions were largely inseparable, whereas the forest–treeline transition explained only a small, though statistically detectable, fraction.

Treeline effects were context-dependent at different sites: shifts in the abundance of individual taxa, pronounced in the Alaska and Brooks Ranges and weak in Interior Alaska; reduced alpha diversity at the drought-limited Interior Alaska treeline; and an increased ECM share of the root-associated community at all three treelines, significant in the Brooks Range. These differences were not explained by tree age or by fine-scale spatial structure. Six of the eight ECM taxa that responded were enriched at the treeline, and plant saprotrophs declined at all three treelines. The most consistent belowground signature of the treeline in this system is therefore a shift in guild balance towards mycorrhizal fungi, visible both in the guild as a whole and in the direction of its responding members.

Neither RAF community composition, nor alpha diversity, nor guild abundance was associated with individual radial growth over the previous 5–30 years, under any of three growth measures and after accounting for site, habitat and tree size. Within the precision of our data, ECM diversity is not a useful predictor of long-term performance in mature *P. glauca*.

## Acknowledgements

The authors thank the student helpers, Katherina Kahlke and Frederik Märker, for their hard work in the field, particularly for digging out and carefully cleaning fine roots under challenging weather conditions. We are also very grateful to our excellent guide, Andreas Burger, for his support in the field. We thank Elke Seeber for her help with organizing the field trip, and Martin Wilmking and members of his working group (especially Timo Pampuch and Melanie Zacharias) for establishing the research plots and sharing valuable information about them. We are grateful to Mo Wirth for the thorough and careful work on DNA extraction from the fine roots. Finally, we thank Ulrich Möbius for his assistance with soil pH measurements and Mario Trouillier for his valuable advice on data analysis.

## Author contributions

KK: Conceptualization, Data curation, Formal analysis, Investigation, Resources, Methodology, Visualization, Writing – original draft

SB: Investigation, Formal analysis, Resources, Writing – original draft MB: Conceptualization, Supervision, Writing – review and editing

MS: Conceptualization, Funding acquisition, Supervision, Writing – review and editing

