## Supplementary material for "Ectomycorrhizal fungi associated with mature white spruce (*Picea glauca*) across three Alaskan treelines": SF1

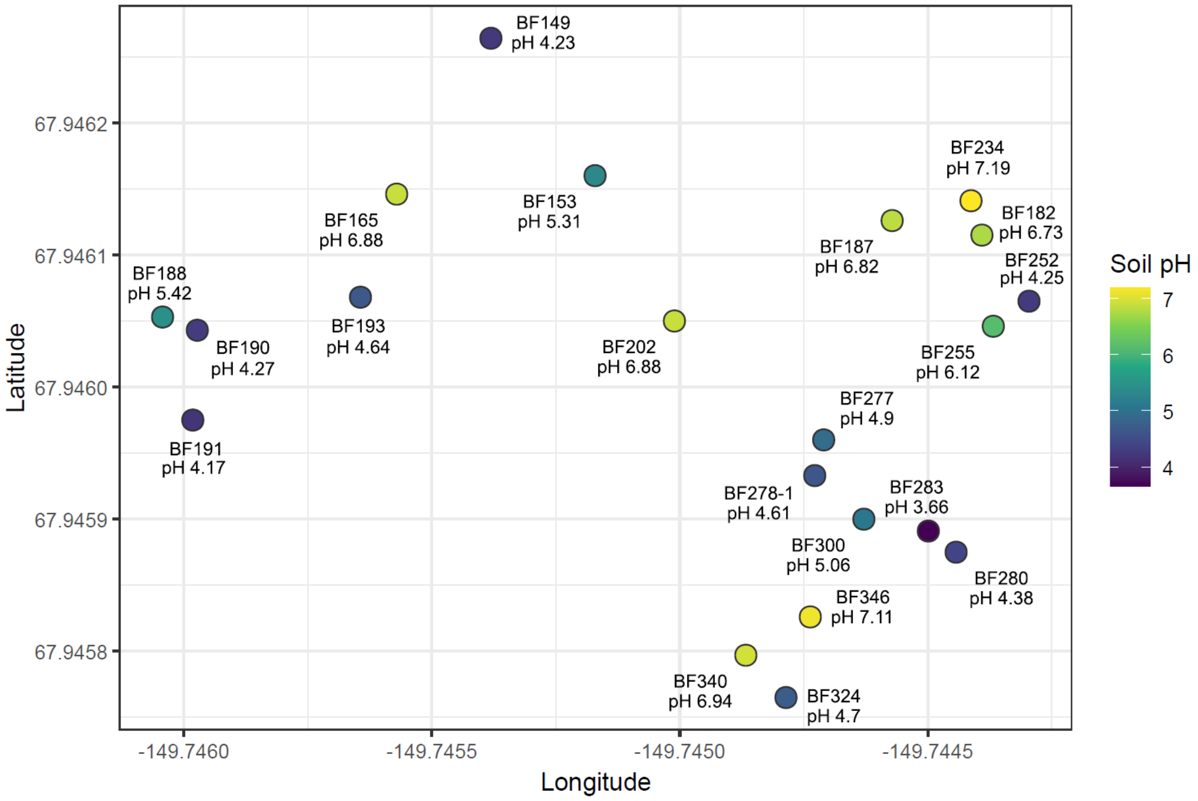


**Supplementary Figure SF1.** Spatial distribution of *P. glauca* trees in the Brooks Range forest and the pH of adjacent soil.
