## Supplementary material for "Ectomycorrhizal fungi associated with mature white spruce (*Picea glauca*) across three Alaskan treelines": SF2

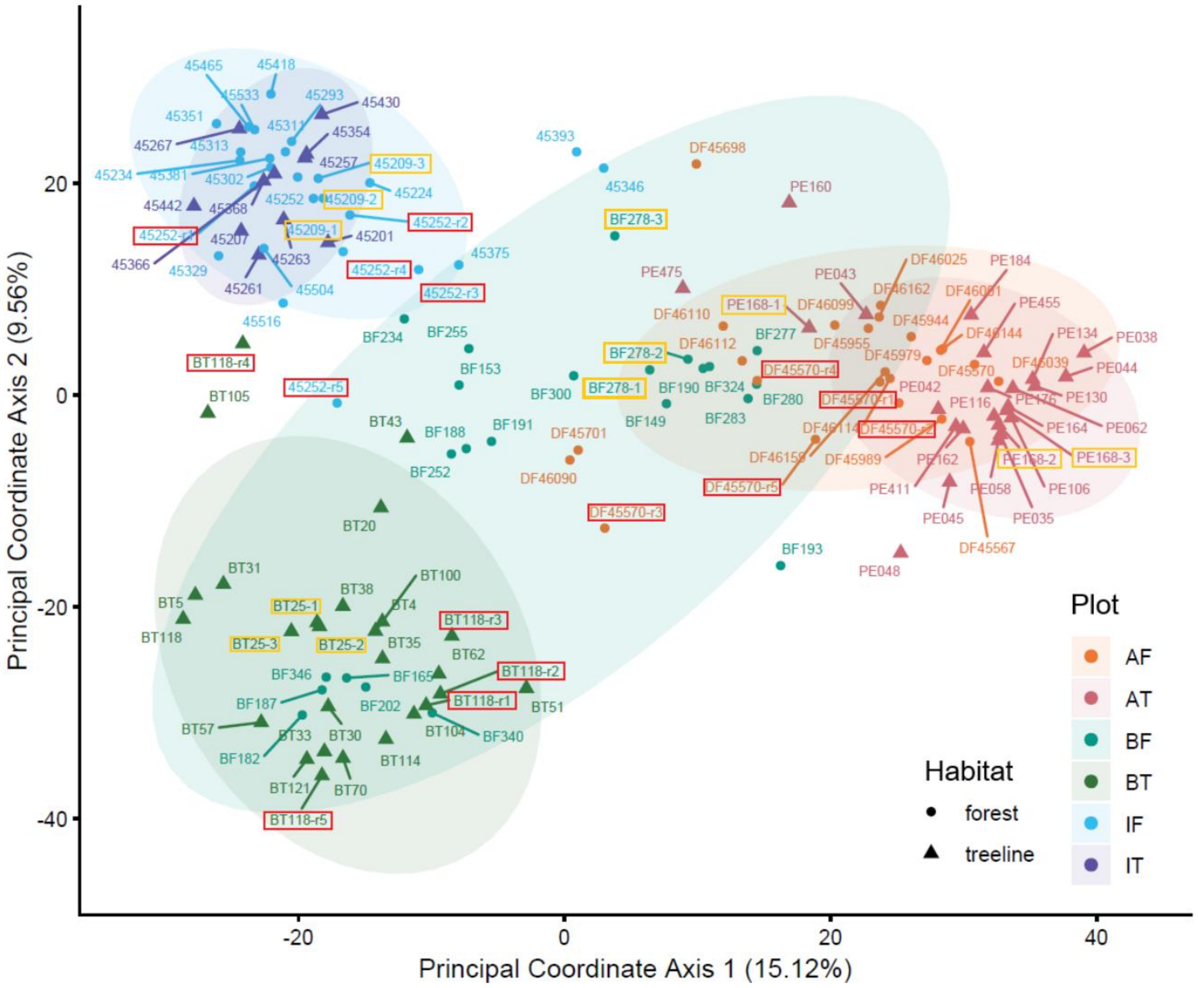


**Supplementary Figure SF2.** Ordination diagrams generated from Euclidean distances via Principal Coordinate Analysis for all 463 OTUs of root-associated fungi. Distances were calculated between normalized counts of OTUs found in pooled root samples of 111 trees of *Picea glauca*, as well as five root replicates for each of three samples and three technical replicates for each of four samples. Six plots were studied: AF – Alaska Range forest, AT – Alaska Range treeline, BF – Brooks Range forest, BT – Brooks Range treeline, IF – Interior Alaska forest, IT – Interior Alaska treeline. Sample labels of the form "[sample]-[1/2/3]" (yellow boxes) indicate three PCR replicates of the same DNA extract, whereas "[sample]-r[1/2/3/4/5]" (red boxes) indicate five different roots from the same tree.
