## Supplementary material for "Ectomycorrhizal fungi associated with mature white spruce (*Picea glauca*) across three Alaskan treelines": SF3

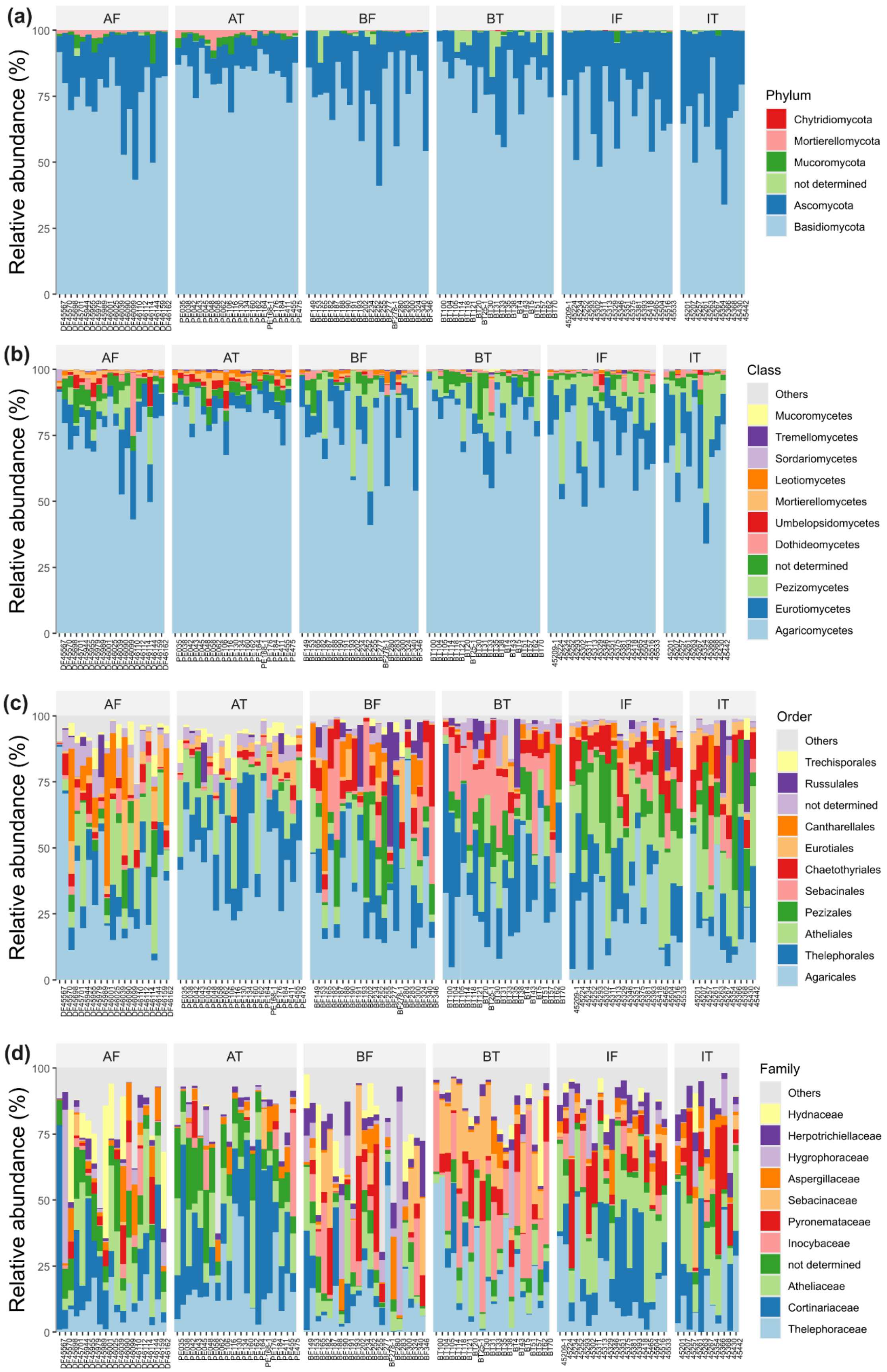


**Supplementary Figure SF3.** Relative abundance of 463 OTUs assigned to (a) phylum, (b) class, (c) order, (d) family found associated with 111 trees of *Picea glauca across six study plots:* AF – Alaska Range forest, AT – Alaska Range treeline, BF – Brooks Range forest, BT – Brooks Range treeline, IF – Interior Alaska forest, IT – Interior Alaska treeline.
