## Supplementary material for "Ectomycorrhizal fungi associated with mature white spruce (*Picea glauca*) across three Alaskan treelines": SF4

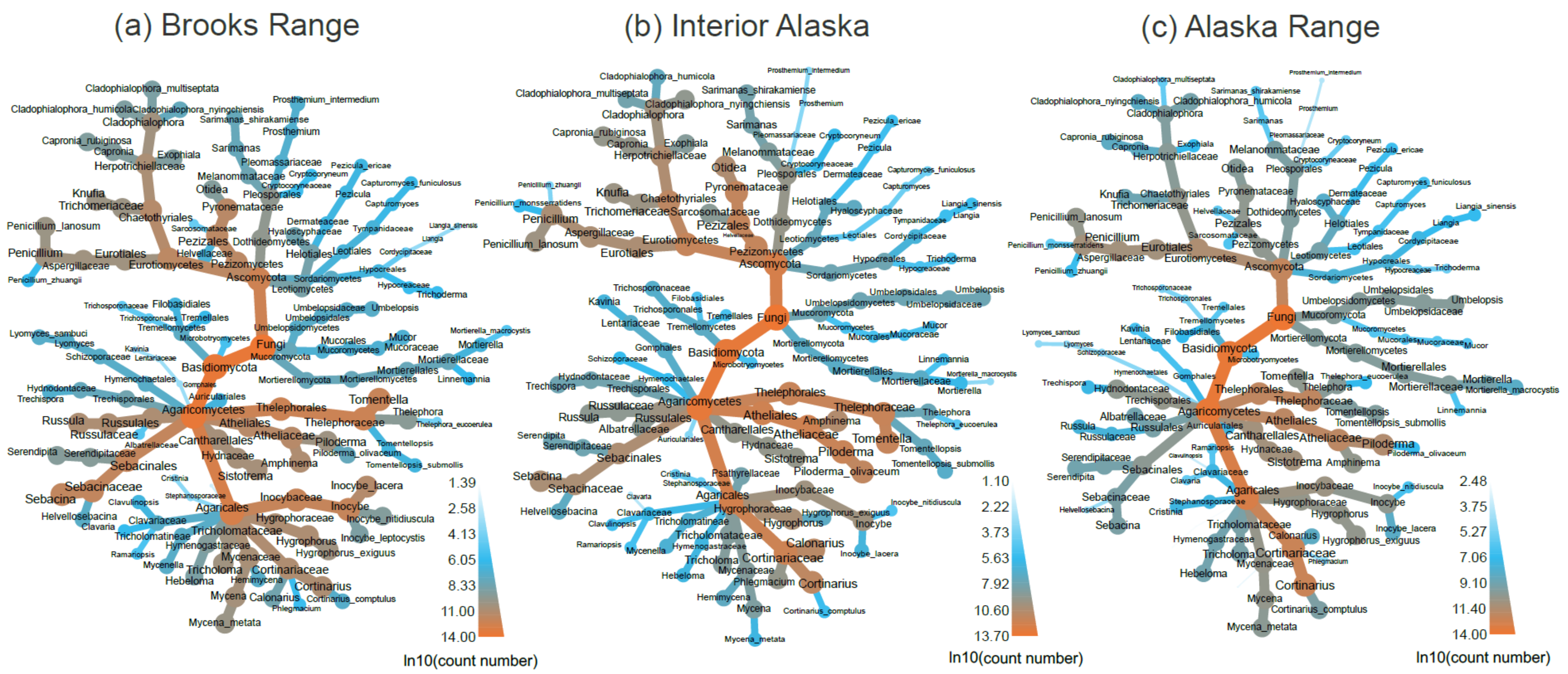


**Supplementary Figure SF4**. Taxonomic heat trees illustrating diversity of fungal taxa identified in three study sites: (a) Brooks Range, (b) Interior Alaska, and (c) Alaska Range. Only taxa represented by at least two OTUs were included. Node size and colour indicate the observed log10-transformed read counts.
