## Supplementary material for "Ectomycorrhizal fungi associated with mature white spruce (*Picea glauca*) across three Alaskan treelines": SF5

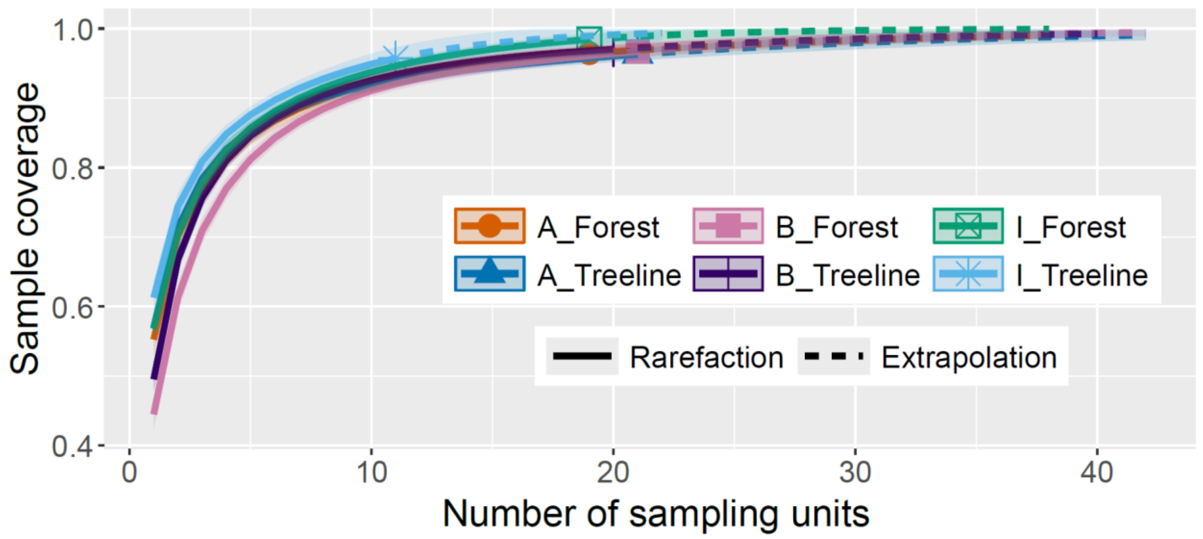


**Supplementary Figure SF5.** Sample-size-based rarefaction and extrapolation curves for forest and treeline trees of *Picea glauca* across six study plots: A – Alaska Range, B – Brooks Range, I – Interior Alaska.
